# The interaction of climate history and evolution impacts alpine biodiversity assembly differently in freshwater and on land

**DOI:** 10.1101/2021.12.17.472935

**Authors:** Luiz Jardim-deQueiroz, Carmela J. Doenz, Florian Altermatt, Roman Alther, Špela Borko, Jakob Brodersen, Martin M. Gossner, Catherine Graham, Blake Matthews, Ian R. McFadden, Loïc Pellissier, Thomas Schmitt, Oliver M. Selz, Soraya Villalba, Lukas Rüber, Niklaus Zimmermann, Ole Seehausen

## Abstract

Quaternary climate fluctuations can affect biodiversity assembly through speciation in two non-mutually-exclusive ways: a glacial species pump, where isolation in glacial refugia accelerates allopatric speciation, and adaptive radiation during ice-free periods. Here we detected biogeographic and genetic signatures associated with both mechanisms in the generation of the European Alps biodiversity. Age distributions of endemic and widespread species within aquatic and terrestrial taxa (amphipods, fishes, amphibians, butterflies and flowering plants) revealed that endemic fish evolved only in lakes, are highly sympatric and mainly of Holocene age, consistent with adaptive radiation. Endemic amphipods are ancient, suggesting preglacial radiation with limited range expansion and local Pleistocene survival, perhaps facilitated by a groundwater-dwelling lifestyle. Terrestrial endemics are mostly of Pleistocene age, and are thus more consistent with the glacial species pump. The lack of evidence for Holocene adaptive radiation in the terrestrial biome may be attributable to a faster range expansion of these taxa after glacial retreats, though fewer stable environments may also have contributed to differences between terrestrial areas and lakes. The high proportion of young, endemic species make the Alps vulnerable to climate change, but the mechanisms and consequences of species loss will likely differ between biomes because of their distinct histories.

## 1. Background

Immigration, speciation and extinction are the three main processes underlying the assembly of biodiversity in island-like habitats [1–4]. The relative contribution of these processes depends on size, isolation and fragmentation of the region, ecosystem or habitat. For instance, immigration rates decrease with increasing isolation, extinction rates decrease with increasing area, and rates of in situ speciation increase with both area, isolation, and fragmentation [1,2,5–7]. The occurrence and interaction of these processes over geological history leave strong imprints in the contemporary structure of regional and local species assemblages, including phylogenetic structure and relatedness. The species age distribution, and the nature and degree of endemism are some of the resulting biodiversity features [8].

Some of the mechanisms that can lead to endemism are through cladogenetic speciation, i.e., when an ancestral species diverges into two or more derived species within an island, archipelago or geographic region, or through anagenetic speciation, when a local or regional population or set of populations diverges from its progenitors outside the island, archipelago or region [3]. Recent cladogenetic and anagenetic speciation both result in neoendemic species, which are young species with geographically restricted distributions. Moreover, if cladogenetic and anagenetic speciation are the main processes behind regional biodiversity assembly, a regional biota can be composed of many relatively young and closely related species. Non-endemic species, in turn, are generally more widespread because they either have immigrated to the focal region from the outside after range expansion or they have arisen in the focal region and had time to spread beyond it. Such non-endemic species are also expected to be older, according to the ‘age and area’ hypothesis [9], which predicts that older species have had more time to disperse and hence become geographically more widespread, whereas younger species often are still confined to smaller ranges. However, the ‘age and area’ hypothesis assumes biome stability (including climatic stability) and does not consider factors other than age that could in fact have strong effects on species range sizes. For instance, population extirpation or local extinction [10], the presence and movement of physical or climatic barriers in space and time [11], changes in habitat size through time, variation between lineages in the ecological versatility and evolutionary adaptability of species [12], variation in species dispersal ability [13,14] and ecological interactions [15] can all be important predictors of species range size in addition to species age. The interaction of these factors can explain, for instance, why some species, despite being so old, are geographically narrowly confined in the present time (geographical relicts or paleoendemics) [16].

Physically rugged mountain landscapes at lower and intermediate latitude, such as those of the Alpine bioregion of Europe (hereafter the European Alps), are hotspots of biodiversity and endemism [17–19]. In such environments, endemism and species radiations arise through the interaction of dispersal limitation with steep ecological gradients and often archipelago-like habitat structures [20–23]. In the European Alps, multiple terrestrial taxa have undergone local radiations leading to the emergence of endemic clades in several groups such as flowering plants [24–26] or butterflies [27,28]. Furthermore, some of the largest endemic radiations in European freshwater habitats also took place in or around the Alps, especially for amphipods [29–32] and fish [33,34]. Importantly, these radiations occurred in habitats that are geographically isolated from similar habitats elsewhere, but surrounded by less isolated habitats, containing diverse assemblages of widely distributed taxa. For example, mountain-tops surrounded by lowlands, or permanently cold, deep lakes isolated from other such lakes by the seasonally relatively warm, shallow flowing water of rivers.

The climatic and habitat instability driven by the Quaternary climate fluctuations [35] has interacted evolutionary and ecological processes to shape biodiversity in the Alps [34,36–41]. This includes influences on speciation, extinction and immigration of lineages, and the reshaping of species abundance, range distribution, richness and genetic diversity patterns [42–44]. An important fraction of biodiversity in the Alps is due to recently colonizing species that immigrated into the region from far away (such as central Asia), or expanded their range from adjacent regions but have not yet speciated in the area. On the other hand, endemic biodiversity may have emerged through two alternative, non-exclusive mechanisms both driven by the succession of glacial–interglacial cycles: (1) the glacial species pump [45,46], and (2) adaptive radiation during interglacial periods (hereafter adaptive radiation) [47] (figure 1). The glacial species pump is a process in which allopatric speciation is accelerated via the isolation of small populations in glacial refugia. It operates when the expansion of glaciers makes large areas of a species’ range unavailable, but leaves isolated pockets of suitable habitat (figure 1b; Hewitt 2000; Hewitt 2004; Holderegger and Thiel-Egenter 2009; April et al. 2013). It can therefore be expected that glacial pump creates assemblages composed of many species that originally emerged in allopatry (figure 1c) but might have come into secondary contact more recently (figure 1e). The role of refugia in promoting species persistence and in glacial vicariant speciation has been widely reported for multiple extant European taxa, both animals and plants [36,37,e.g. 50–53].

**Figure 1.**
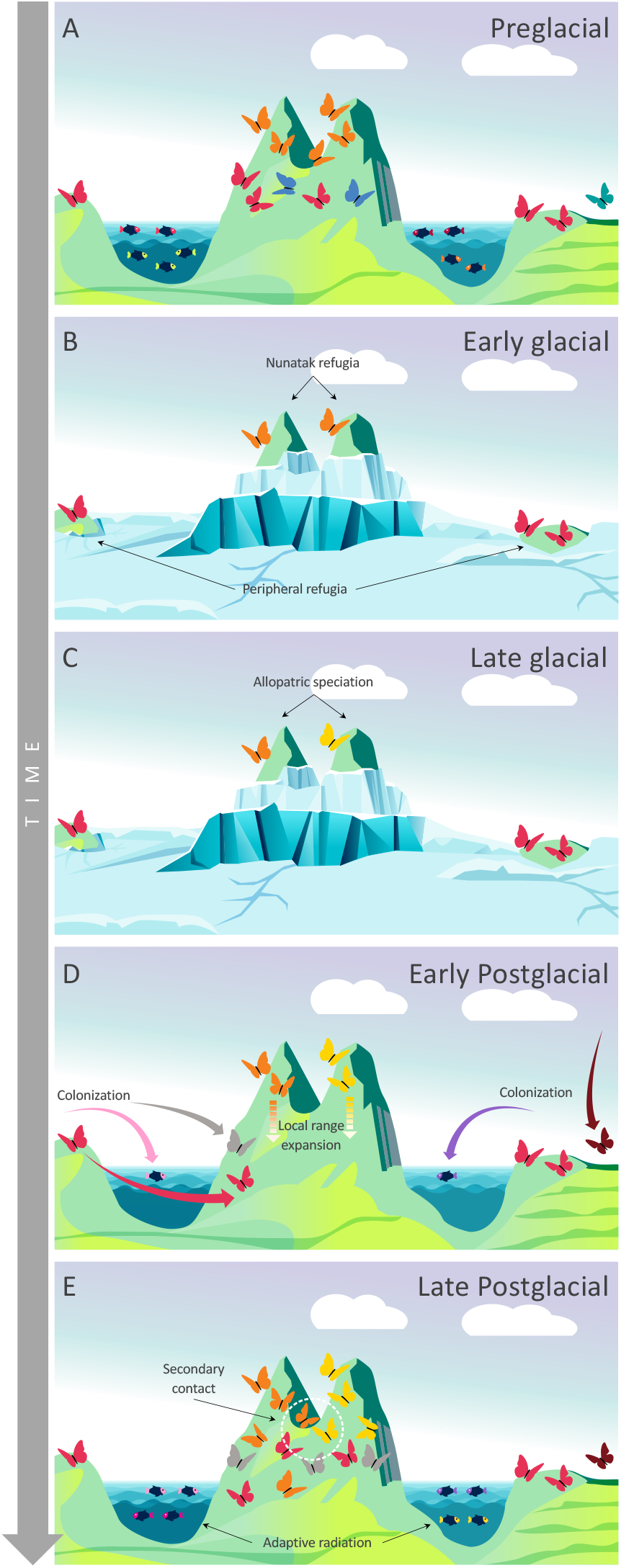
Evolutionary and ecological history of a hypothetical biodiversity assembly in an alpine-like system. A) Biodiversity in a preglacial phase. B) Early glacial phase: glacial periods erase freshwater habitats and fragment the terrestrial biome. Some populations survive in refugia and C) can diverge into distinct species through allopatric speciation. D) The retreat of glaciers opens up new, unoccupied habitats offering ecological opportunities for colonizers. E) Some colonizers undergo adaptive radiation and niche space is filled up again.

After each glacial maximum, the retreat of glaciers opens up new, unoccupied habitat in both terrestrial and aquatic ecosystems, which may lead early colonists to diversify *in situ* via adaptive radiations [54] (figure 1b). Adaptive radiation takes place when an ancestor diversifies ecologically and phenotypically, giving rise in a relatively short period of time, to multiple species that are ecologically distinct [55]. Emblematic examples of adaptive radiation during inter- or postglacial periods are associated the emergence of many lakes in the Holarctic realm by the end of the last glacial period, providing opportunities for many radiations of freshwater fish that occurred in parallel [33,34,47,56–59]. Such process would be expected to generate sympatric or parapatric species by *in situ* cladogenetic speciation [23]. In addition, postglacial expansion of populations also has the potential to bring together lineages that had previously diverged in Pleistocene refugia [60]. Secondary contact of lineages can also be important in adaptive radiation, either through causing ecological character displacement in sympatry occurring in response to competition [55] (Schluter 2000, or through the occurrence of hybridization which may facilitate the onset of adaptive radiations upon colonization of new environments [61,62]. This ‘hybrid swarm origin’ hypothesis of adaptive radiation posits that functional genetic variation, which becomes enriched in hybrid zones, elevates the evolvability and response to natural selection in hybrid populations [61,63].

Despite its large information content for investigating hypothesis regarding biodiversity assembly and to understand biogeographic patterns in species distributions [64], few studies have made use of the species age distribution (SAD) [41,65,66], and fewer had investigated SAD from a multi-taxon perspective [8,67]. We conducted a comparative phylogenetic analysis to quantify SADs and the extent and type of endemism in aquatic and terrestrial ecosystems of the European Alps. Our work focuses on several taxonomic groups: amphipods, fish, butterflies and amphibians with nearly complete taxon sampling, as well as 15 nearly completely sampled representative genera of perennial flowering angiosperms plants (henceforth “plants”). We predicted that SADs support a scenario with dominance of the glacial species pump for the origin of endemism in terrestrial groups, with species dating to the Pleistocene; whereas SADs may indicate a prominent role of postglacial adaptive radiation for groups that depend on open water habitats, i.e., fish. This is because high altitude ranges in the terrestrial habitats became fragmented but not completely erased during glacial maxima, whereas year-round open water bodies as habitat for fish were entirely absent during glacial maxima, with a few exceptions at the edge of the southern Alps [68]. We also predicted that SADs of both amphipods and amphibians will resemble fully terrestrial taxa more than those of fish. Amphipods occupy both open and subterranean freshwater habitats, and some of the subterranean habitats persisted during the glacial maxima [69], and most amphibians require open water bodies only during spring and summer and are terrestrial for the remainder of the year, while some lack an aquatic life stage altogether [70]. Therefore, some amphibian species likely found Pleistocene refugia within the Alpine region during the glacial maxima [71,72]. Regarding non-endemic species, we did not expect large differences in age structure among taxa because non-endemics tend to be widespread, are probably mostly older and diversified on a wider geographical scale drive by processes tha may have been decoupled from the climate dynamics of the Alps. Therefore, we predicted that non-endemic species are older on average than endemics. Because dispersal ability, distance between sink (new habitats that become available after glacial maxima) and sources of colonization, historical stability and heterogeneity of habitat occupied vary between the taxonomic groups, the magnitude of the difference in species ages between non-endemic and endemic species will be taxon-dependent.

## 2. Methods

Our work focuses on the European Alps, following previous delimitations of European high mountain systems [73] but including the peripheral lowland areas, where most of the perialpine glacial lakes are located. To select the taxonomic groups to be included in this work, we looked for lineages with reliable distribution data, robust dated phylogenetic trees that include most of the diversity of the given lineage, and/or genetic data for most of the recognized species, so that we could estimate and calibrate phylogenetic trees where none existed. Then, we chose five major taxonomic groups to represent the terrestrial and aquatic alpine and pre-alpine biomes of the region: nearly all known regional species of amphipods, fishes, amphibians and butterflies, and 14 nearly completely sampled clades of flowering plants (*Homogyne, Petasites, Tussilago, Campanula, Jasione, Phyoplexis, Knautia, Androsace, Primula, Soldanella, Gentiana, Saxifraga, Carex and Festuca*). Only species native to the European Alps were considered. To assemble species age distributions (stem age, i.e., time since divergence from closest relative in million years, Ma), we combined published time-trees and our own estimates (electronic Supplementary Methods).

### (a) Endemism and speciation mode

Species were considered endemic if they naturally occur only in the alpine and/or perialpine regions of the Alps. When a species was classified as endemic, we assigned its speciation mode to anagenetic speciation (Alpine species diverged from its non-Alpine sister species, but did not undergo *in situ* diversification) or cladogenetic speciation (Alpine species emerged through *in situ* diversification, with either the non-endemic sister being native to the region too, or the two or more sister species all being Alpine endemics), following Rosindell and Phillimore (2011) (electronic supplementary material, tables S1). To apply these definitions, we first identified the position of each of the endemic species within the phylogeny (electronic supplementary material, file S2). Then, if the endemic species was nested within a clade composed mainly of species that occur outside of the Alps, and its direct sister species occurred only outside the Alps, we assumed anagenetic speciation. If the species was nested within a group mostly of species native to the Alps, we assumed cladogenetic speciation.

### (b) Comparisons of species age distributions

We performed multiple permutation tests on the distribution of age estimates assembled from all the species to identify differences in the age distributions among and within major clades. Here, we asked 1) whether age distributions differed between endemic and non-endemic species, overall and within each taxonomic group independently; 2) whether species age distributions differed among taxonomic groups; 3) whether non-endemic species age distributions differed among groups; and 4) whether endemic species age distributions differed among groups. When a difference was significant, if a significant result was found, we performed post-hoc pairwise permutations to identify which distribution and taxonomic group (or groups) were distinct from one another, using Bonferroni correction for each group of analyses. These analyses were performed using the function ‘oneway_test’ of the library ‘coin’ [74] in R 4.0.2 1106 [75] with 10,000 resamples with a distribution approximated via Monte Carlo resampling.

## 3. Results

A total of 617 species were included in our analyses: 39 amphipods, 124 fishes, 31 amphibians, 245 butterfly and 178 plant species (electronic supplementary material, table S1). Approximately half of all fish and amphipod species were found to be endemic to the European Alps (45.2 % and 49 %, respectively), whereas smaller fractions of 12 %, 13 % and 30 % of the butterflies, amphibians and plants, respectively, were found to be endemic (figure 2). Our analysis of speciation mode suggests that approximately half of the endemic amphipods and plants, and one third of butterfly species emerged by cladogenesis (53 %, 52 % and 36 %, respectively). Cladogenesis was also inferred to be the mode of speciation for the majority of the endemic amphibian and fish species (75 % and 98 %, respectively) (figure 2).

**Figure 2.**
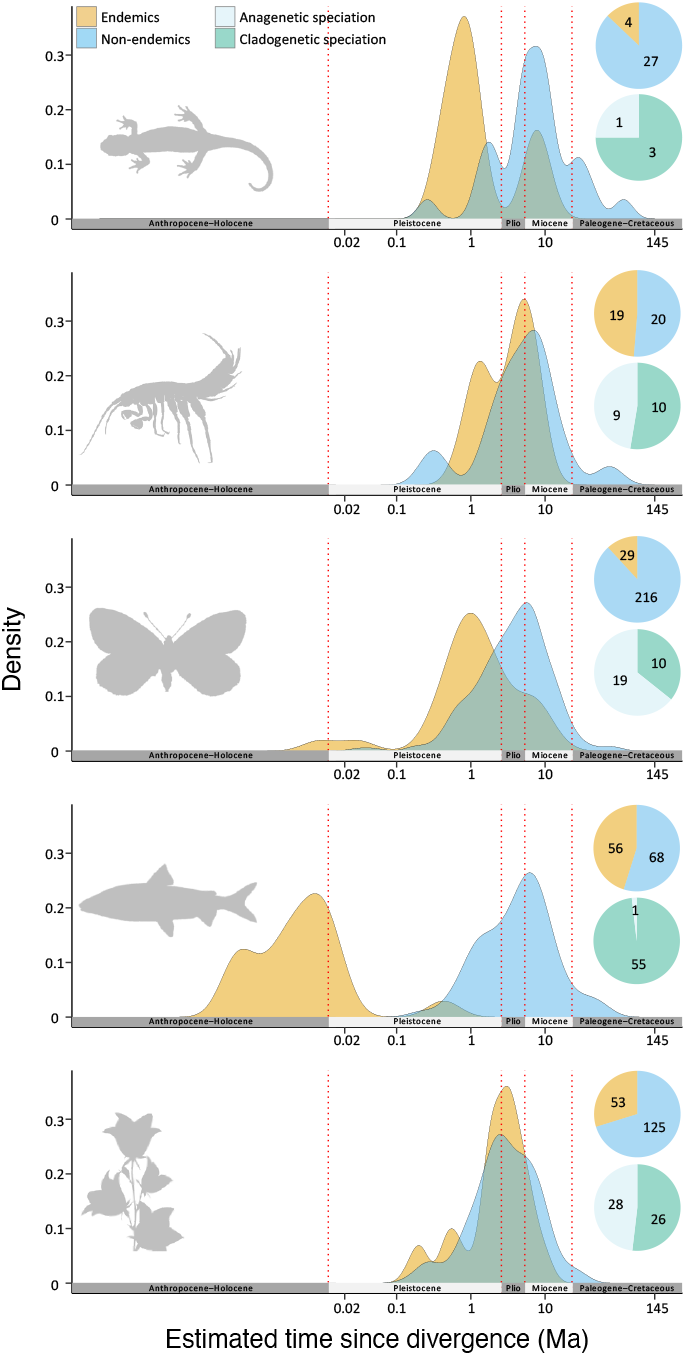
Species age distribution (SAD) of endemic and non-endemic species of, from top to bottom, amphibians, amphipods, butterflies, fish and flowering plants. Pie charts show the proportion of endemic and non-endemic species as well as the proportion of endemic species that have emerged through cladogenetic or anagenetic speciation.

We found that 94 % of the extant species that now occupy the Alps, irrespective of endemism status, emerged over the past 15 million years (from the Middle Miocene) (figure 2). Endemics were overall younger than non-endemics (p-value < 0.0001), which was also true for each taxonomic group when analysed individually (electronic supplementary material, table S9). We found the species age distributions among non-endemic species to be similar between taxonomic groups (figure 2), with most of the species ages spanning the Late Pleistocene to Early Miocene (90 % fell between 0.3 and 20 Mya). The only exception was the comparison of plants *vs*. amphibians, as we found non-endemic amphibians older than non-endemic plants (p < 0.0001; electronic supplementary material S10). SADs of endemic species were also similar among taxa (90 % fell between 0.25 and 8 Ma), except for fish, which are younger than any other group (90 % fell between 0.6 and 114 Ky; p < 0.0001; figure 2; electronic supplementary material S11).

We observed the following patterns in the SAD: i) endemics are younger than non-endemics in all taxonomic groups, and ii) there is a high degree of similarity among taxa in the age distributions when endemics and non-endemics are analysed separately. However, we found on one hand that while most endemic fish species arose in the Holocene through cladogenesis, very few endemic butterflies and no endemic species in the other groups arose in the Holocene. On the other hand, only very few endemic fish (all from the southern periphery of the Alps) date to the Pleistocene whereas most of the endemic butterflies and many other endemic species date do.

## 4. Discussion

Inferring past evolutionary process from the structure of current biodiversity is one of the goals of macroecology, macroevolution and biogeography. Species age distributions among regional biota is one informative aspect of this structure [64]. We combined estimates of species origination times with information on endemism and mode of speciation (cladogenetic vs. anagenetic) to investigate the dominant mechanisms of biodiversity assembly in different taxa and major biomes of the European Alps. We found that most of the species diversity in fish, amphipods, amphibians, butterflies and plants is relatively young, emerging at the beginning in the Middle Miocene, which coincided with the period of maximum geological uplift in the region that culminated with the formation of the Alps [76].

Our multi-taxon comparative analysis had the advantage of directly comparing independently evolving and ecologically distinct clades in the same region. It showed that speciation timing was dramatically different between terrestrial taxa and those aquatic taxa that require permanent open surface water (i.e., fish). While 80 % of the Alp’s endemics in the terrestrial groups originated between the Late Miocene and Late Pleistocene, most endemic fish species arose only after the final retreat of the glaciers and re-establishment of permanent open water bodies in the formerly glaciated areas. Combined with the observation that the vast majority of endemic fish are products of cladogenetic speciation, this suggests that the assembly process of the fish fauna of the Alps is dominated by an interaction between colonization from outside the region and adaptive radiation during the last postglacial period, but similar processes may have cyclically repeated themselves in previous interglacial periods, alternating with extinction during glacial maxima. In contrast, for the terrestrial groups, our results suggest that colonization from outside the region and the glacial species pump are the dominant mechanisms, as anagenetic speciation was more important in these taxa, and endemic richness assembled throughout the Pleistocene. Interestingly, we observed some postglacial speciation in butterflies, coinciding with the major mode in fish, but being dwarfed in butterflies by the much larger Pleistocene mode. We suggest that general, non-exclusive mechanisms underlay these contrasting patterns: 1) Quaternary climate fluctuations that accelerate allopatric speciation during cold stages but open up new ecological opportunities for adaptive radiation during interglacial periods; 2) variation among groups in their dispersal ability and associated rate of range expansion, and finally 3) the influence of variation in seasonal and inter-annual habitat stability in either constraining or promoting adaptive radiation. Below we discuss each of these mechanisms and how they may have affected diversification in the different taxa studied here.

### (a) The role of Quaternary climate fluctuations

We suggest our finding that most endemic fish are of postglacial origin, while endemics in other groups arose in the Pleistocene or earlier, is explained by the different effects that the Quaternary climate oscillations had on freshwater versus terrestrial habitat [68]. Permanent open surface water habitats, as required by fishes, were absent in the glaciated parts of the Alps during the Last Glacial Maximum (LGM), because all lakes and river valleys were covered by thick glaciers. Therefore, the complete lack of endemic fish species older than 20,000 years on the northern and western flanks of the Alps is likely due to local extirpation of all fish populations across the region during the LGM. With the progressive retreat of glaciers, which achieved their modern configuration in the Late Holocene, fish would then have returned to the region from areas located in downstream sections of the large rivers, that were often far from the alpine region especially on the North face of the Alps [e.g. 77]. This Holocene recolonization by older widespread species explains the large fraction of old, widespread and non-endemic fish species in the northern and western Alpine region.

The first fish to colonize after the LGM were probably species adapted to cold water conditions, such as salmonids und sculpins, that would have lived nearby in the rivers of the Pleistocene tundra downstream of the Alpine glacier shield. Salmonids are indeed known for their remarkable colonizing ability and rapid establishment in postglacial freshwater habitats [78]. These fish would have encountered ecological opportunities in the emerging large and deep lakes of the region and radiated into many distinct species as they adapted to the vacant niches associated with distinct lacustrine zones. This process likely generated the young endemic species, nowadays predominantly in three lineages, whitefish (*Coregonus* [79]),, chars (*Salvelinus* [80]) and sculpins (*Cottus* [34]), that have rapidly radiated in perialpine lakes. The very few old, relic endemic fish species in the region that date to prior to the Holocene, are the lake herring *Alosa agone*, and two trouts of the genus *Salmo* (*S. carpio* and *S*. sp. ‘Blackspot’). These three species are endemic to lakes in Northern Italy and southern Switzerland, a region where probably not all lakes were fully covered by ice sheets during the LGM [68]. Therefore, these species likely originated during earlier interglacials, when southern perialpine lakes would have become extensive, and then found refugia during the LGM to persist to the present day [81]. That there are no young postglacial species among the non-endemic fish is perhaps due to insufficient time and connectivity between lakes to allow new species to arose in deep lakes elsewhere in Europe (e.g., in northern Germany and Scandinavia) in the Holocene prior to expanding their range into the Alps or vice-versa. It is important to mention that adaptive radiation of fish are far more frequent in deep lakes [82,83], while riverine adaptive radiations are rare (but see [84–86]).

Unlike for fish, our analysis revealed that all endemic (and all nonendemic) amphipods, the second fully aquatic taxon in our data, emerged during or before the Pleistocene. This could be because amphipods can persist in smaller water bodies than fish, and many species in this group are ice-associated, being able to survive under ice cover and in its immediate forefield [87]. Both factors would have allowed some species to persist in the region throughout the glaciations. Additionally, as suggested for other invertebrates [69], some species likely survived in subterranean refugia, such as caves or groundwaters, habitats occupied by many freshwater amphipod species today, notably species of *Niphargus*, the most species-rich group in the region [29,32].

We found that endemic species in the terrestrial groups are much older than endemic fish species. Pleistocene refugia are hypothesized for terrestrial taxa in many geographically restricted areas on nunataks, i.e, mountain peaks that have never been glaciated [88], or surrounding the Alps [41,89–92], such that extinctions during the glacial cycles did not wipe out the terrestrial fauna entirely. It is very likely that terrestrial organisms, particularly butterflies [41,93], plants [94–96] and amphibians [71,72] survived in glacial refugia in the Alps and at its periphery. Therefore, differential impacts of Quaternary climate fluctuations and the resulting glaciations on different habitats and taxa go a long way helping to explain extant patterns of diversity and endemism in the region.

### (b) Dispersal ability to explain postglacial radiation

Dispersal ability often negatively correlates with rapid niche evolution. The evolutionary response to local environmental and ecological conditions tends to be faster in taxa with limited dispersal, leading to faster niche shifts and higher rates of speciation and adaptive radiation [97,98]. Therefore, the intriguingly few cases of postglacial speciation in fully terrestrial species and amphibians could be related to the dispersal rates imposed by the environments they occupy. Terrestrial taxa experience, in general, less dispersal limitation than freshwater taxa. For example, many species have acquired adaptations for aerial dispersal, such as active flying in butterflies [99,100] or passive airborne propagation in many plants [101], allowing such taxa to disperse virtually in all directions. Conversely, freshwater-bound taxa need to navigate the dendritic landscapes of rivers and lakes to disperse, making it a lot more difficult to reach isolated habitat patches [102,103]. Given these limitations to dispersal for many freshwater taxa, postglacial dispersal may have happened at a much slower pace in fish than for most terrestrial taxa. Terrestrial species may, hence, have expanded their ranges faster after glacial retreat, also likely facilitated by the proximity of refugia to the Alps (including inner-alpine refugia), resulting in faster recolonization of the newly open landscape through long-distance dispersal and range expansion. Some recent studies have shown that many plant species rapidly and substantially expanded their range during the recent postglacial period [104–106]. Faster filling of emerging terrestrial habitats through range expansion left fewer opportunities/less time for the first colonists to undergo ecological speciation and adaptive radiation in response to ecological opportunity among terrestrial groups than among aquatic taxa. To test the relative importance of dispersal limitation versus other aquatic/terrestrial differences, future work could investigate mainly aquatic taxa with strong aerial dispersal abilities, such as Odonata and other insects that spent most of their life cycle in freshwater and have short but highly dispersive terrestrial adult phases.

### (c) Seasonal and interannual environmental variation limiting ecological speciation

Habitat stability in the Postglacial Era may have been an additional factor explaining the larger number of Holocene speciation events in fish, but not in terrestrial groups in our study. Theory and models suggest that environmental fluctuations and stochasticity can reduce or even inhibit ecological speciation in unstable habitats [107,108]. Rapid variation in environmental conditions, both seasonal and interannual, make adaptation difficult and ecological speciation nearly impossible.

Environmental conditions in terrestrial ecosystems are much more variable than aquatic ecosystems [109,110], especially large and deep lakes, both in terms of seasonal and year-to-year variation. For instance, whereas seasonal variations in solar irradiance, temperature and snow cover make the high mountain terrestrial habitat extremely seasonal, with large year-to-year variation in the onset and duration of seasons [111,112], they are nearly constant through the year in the deeper parts of lakes [113,114]. The longer growing and reproductive season, despite low productivity, and the much more stable environment in deep lakes may create increased opportunities for ecological speciation and adaptive radiation compared to the alpine terrestrial ecosystems.

In addition, despite their greater temporal stability, deep lakes also have much steeper environmental gradients because pressure, light and temperature all change much faster with depth in water than with elevation in the terrestrial realm. This unique property of water may explain the high frequency of ecological speciation in deep lakes, with sister-species being spatially very close to each other but occupying different water depths [58], as observed among East African cichlids [115] and Alpine whitefish [116].

## 5. Final considerations

We suggest that the formation of the unique biota of the European Alps was driven by interacting mechanisms: non-random Pleistocene survival, postglacial immigration, vicariant speciation during glacial maxima and adaptive radiation in the Postglacial. These interacting mechanisms left distinct imprints on the age structure of regional assemblages in different biomes and associated taxon groups. Historical factors (Quaternary climate fluctuations and Pleistocene refuge availability) impacted freshwater and terrestrial biomes in different ways, and contemporary ecological factors such as environmental stochasticity and dispersal limitations also vary between these biomes, shaping them very differently through ecological and evolutionary processes. *In situ* speciation and adaptive radiation were prominent in fish, but occurred mainly after the LGM, and only in deep lakes, likely due to the unavailability of suitable freshwater habitat during the LGM and the stable conditions within habitats after the LGM. Amphipods and all terrestrial clades have much older endemic species, perhaps because their ecology (i.e., cold-resistant and groundwater-dwelling in amphipods) and the availability of Pleistocene refugia within the region allowed many species to survive the LGM. At the same time, none of the terrestrial groups generated many young postglacial species, likely because higher Pleistocene survival and faster postglacial niche filling through range expansion left fewer ecological opportunities and because larger seasonal variation in the terrestrial environment places constraints on ecological speciation.

Knowing the history of biodiversity formation is crucial to establish effective strategies of conservation [117]. For the Alps we show a high fraction of endemism in many groups, with endemic species having survived in some taxa and ecosystems through repeated glacial cycles, while those in others are due to prolific speciation after the retreat of the glaciers. These results improve our understanding of how the Alpine hotspot of species diversity and endemism emerged, and they reinforce that biodiversity in this region is fragile. Endemic species are often range-restricted, show limited population size and are hence much more vulnerable to climate change and other environmental changes than non-endemic species [118], and because of that, they are of high concern for conservation. Even a comparatively small and transient disturbance of an ecosystem can lead to the extinction of young species that evolved in adaptation to specific ecological conditions as has already been observed in the recent past for adaptive radiations of fish in Swiss lakes [116]. The sharp increase of extinction rates driven by human activity in the Anthropocene threatens the biodiversity of the European Alps, and especially that of endemic species [119]. Therefore, this region deserves greater attention to conserve both the regional biodiversity, as well as the eco-evolutionary processes that gave rise to it and that are required to continue operating if biodiversity is to be maintained.

## Acknowledgements

We would like to thank Salome Mwaiko and Rosi Siber for laboratory and GIS support, respectively, and Timothy Alexander for his work in the execution of the *Projet Lac*. Info Fauna provided distribution data of Swiss fishes. For sharing published phylogenetic trees, we thank Natalia Tkach, Adrien Favre, Józef Mitka, Carmen Benitez-Benitez, Markus Dillenberger, Andrew Crowl and Boz□Frajman. We are also grateful to Kay Lucek, Hanna ten Brink and Catalina Chaparro for thoughtful and constructive discussion. Funding from the ETH Board through the Blue-Green Biodiversity (BGB) Initiative (BGB 2020) is acknowledged. ŠB was supported by Slovenian Research Agency through funding programme P1-0184, project J1-2464 and PhD grant.

